# Release of endogenous dynorphin opioids in the prefrontal cortex disrupts cognition

**DOI:** 10.1101/2021.06.27.450102

**Authors:** Antony D. Abraham, Sanne M. Casello, Selena S. Schattauer, Brenden A. Wong, Grace O. Mizuno, Karan Mahe, Lin Tian, Benjamin B. Land, Charles Chavkin

## Abstract

Following repeated opioid use, some dependent individuals experience persistent cognitive deficits that contribute to relapse of drug-taking behaviors, and one component of this response may be mediated by the endogenous dynorphin/kappa opioid system in neocortex. In mice, we find that acute morphine withdrawal evokes dynorphin release in the medial prefrontal cortex (PFC) and disrupts cognitive function by activation of local kappa opioid receptors (KORs). Immunohistochemical analyses using a phospho-KOR antibody confirmed that both withdrawal-induced and optically evoked dynorphin release activated KOR in PFC. Using a genetically encoded sensor based on inert KOR (kLight1.2a), we revealed the *in vivo* dynamics of endogenous dynorphin release in the PFC. Local activation of KOR in PFC produced multi-phasic disruptions of memory processing in an operant delayed alternation behavioral task, which manifest as reductions in response number and accuracy during early and late phases of an operant session. Local pretreatment in PFC with the selective KOR antagonist norbinaltorphimine (norBNI) blocked the disruptive effect of systemic KOR activation during both early and late phases of the session. The early, but not late phase disruption was blocked by viral excision of PFC KORs, suggesting an anatomically dissociable contribution of pre- and postsynaptic KORs. Naloxone-precipitated withdrawal in morphine-dependent mice or optical stimulation of *pdyn*^Cre^ neurons using Channelrhodopsin-2 (ChR2) disrupted delayed alternation performance, and the dynorphin-induced effect was blocked by local norBNI. Our findings describe a mechanism for control of cortical function during opioid dependence and suggest that KOR antagonism could promote abstinence.

## Introduction

Cognitive dysfunction is a common feature of substance use disorders and other psychiatric illnesses [1]. In these clinical syndromes, stress-induced changes in cortical function can exacerbate vulnerability to behavioral disruptions evident during substance use [2]. The neurochemical basis of these responses is not known, but stress activates multiple brain systems that lead to the release of neuropeptide transmitters, including endogenous dynorphin opioids that activate G*α*_i_ protein-coupled kappa opioid receptors (KORs) [3, 4]. Selective KOR agonists, like those in the hallucinogenic sage *Salvia divinorum*, are psychotomimetic in humans [5] and functional polymorphisms in the prodynorphin gene have been correlated with disrupted reversal learning in humans [6], however little is known about the role of the dynorphin/KOR system in physiological control of human cognition.

Stress caused by chronic exposure to addictive drugs increases prodynorphin mRNA expression in brain [7] and the dysphoria experienced during drug abstinence has dynorphin-mediated components [8]. Binge psychostimulant use produces adaptations similar to stress in neural circuits involved with reward processing and reinforcement learning [9, 10]. Chronic exposure to drugs of abuse (e.g. psychostimulants, nicotine, ethanol) increases prodynorphin mRNA in the dorsal and ventral striatum in rodents and humans [7, 11, 12]. Tissue from humans with alcohol use disorders has shown increased dynorphin/KOR mRNA and transcriptional markers in dorsolateral prefrontal cortex, orbitofrontal cortex, and hippocampus [13], suggesting a significant role for dynorphin in cognitive dysfunction following chronic ethanol use [14].

Although the psychopathology of substance use disorders is complex, dynorphins and KOR are expressed in the mammalian cortex [15, 16] and analysis of their role in mouse models of cognition will likely provide useful insights for developing treatments for cognitive dysfunction. In the present study, we demonstrate that pharmacological treatment with a KOR agonist, optogenetic stimulation of PFC prodynorphin mRNA-expressing neurons, or precipitated morphine withdrawal can each significantly increase KOR phosphorylation in the PFC of mice. We then reveal the time course of dynorphin release during precipitated opioid withdrawal in the prefrontal cortex using a genetically encoded fluorescent sensor for endogenous dynorphin in the synaptic space. Cortical regulation of working memory can be readily measured in mice using a delayed alternation procedure [17] and in the present study, we assessed mouse performance in a food-reinforced operant delayed alternation task. We find that pharmacological activation of KOR expressed in PFC as well as endogenous dynorphin released by either naloxone-precipitated morphine withdrawal or optical stimulation of PFC dynorphin neurons disrupted delayed alternation performance. Together, these studies demonstrate that local actions of the dynorphin/KOR system in the prefrontal cortex disrupt cognition.

## Materials and Methods

### Subjects

Adult male C57BL/6 mice ranging from 2-10 months of age were used in these experiments. All experimental procedures were approved by the University of Washington Institutional Animal Use and Care Committee and were conducted in accordance with National Institutes of Health (NIH) “Principles of Laboratory Animal Care” (NIH Publication No. 86-23, revised 1985). Mice were group housed (2-5 mice per cage) and food-restricted to approximately 90% of *ad libitum* body weight when included in the delayed alternation assay. Water was freely available at all times in their home cages. All testing was conducted during the light phase of the 12-h light/dark cycle. The daily food allotment was ~2-3g standard Purina 5053 rodent diet chow per mouse, in addition to food pellets (Bio-Serv Dustless Precision Pellet, Rodent, Grain-Based, 20 mg) consumed during the operant behavioral sessions. We have previously reported estrogen-regulated sex differences in intracellular signaling pathways that alter female responses to KOR agonists and antagonists [18-20]. We observed that the KOR antagonist norBNI is not consistently long-lasting in females, confounding the use of this and other KOR ligands in these studies. For these reasons, we chose to focus these experiments on male mice. Floxed KOR (KOR^lox^) mice were generated by the Institut Clinique de la Souris, in which exon 1 of KOR was flanked by loxP sites [21] and prodynorphin-IRES-Cre (Pdyn^Cre^) mice [22] were obtained from Jackson Laboratories (Jackson Strain 027958).

### Drugs

Racemic (±) U50488 hydrochloride (U50488), norbinaltorphimine dihydrochloride (norBNI), and morphine sulfate were provided by the National Institute of Drug Abuse Drug Supply Program (Bethesda, MD) and naloxone hydrochloride dihydrate was acquired from (Sigma-Aldrich, St. Louis, MO, USA). U50488 (1, 5, and 10 mg/kg), norBNI (10 mg/kg), morphine (10 mg/kg), or naloxone (1 and 10 mg/kg) were dissolved in saline and administered intraperitoneally (IP) in a volume of 10 mL/kg. norBNI (2.5 μg/μL) was dissolved in sterile artificial cerebrospinal fluid (ACSF) and intracranially microinjected. Dynorphin B (Tocris, Bristol, UK) was dissolved in ethanol.

Detailed methods are described in the Supplemental Information section.

### Statistical analysis

All data are presented as mean ±s.e.m. Our sample sizes for behavioral studies are based on those used in Abraham et al. [23] for KOR agonist actions on operant behaviors. Individual data points are shown when possible, and data distribution was assumed to be normal, but not formally tested. Mice were removed from analyses if no viral expression was found or if cannula placement was far from expression site (2 kLight mice excluded). We used two-tailed t-tests, one-way and two-way ANOVA (incorporating repeated measures (paired t-test or RM-ANOVA) where appropriate), and performed Sidak’s, Dunnet’s, or Fisher’s post-hoc tests, specified in text. Mixed effects ANOVA were used when individual mice did not respond during particular time bins to account for missing values. Operant behavior data was collected through a custom-built MED-PC program. Data were analyzed using Prism 9.0 (GraphPad Software; San Diego, California, USA) and custom-built MATLAB software (MathWorks Inc.; Natick, Massachusetts, USA).

## Results

### Stimuli leading to kappa opioid receptor activation in the prefrontal cortex

The dynorphin neuropeptides are expressed by neurons in the prefrontal cortex [24], but physiological and behavioral stimuli necessary to evoke dynorphin release in the cortex have not been identified. To address this question, we used a previously characterized phospho-selective antibody (KORp) [25] to detect agonist-activated KOR that has been phosphorylated by G protein Receptor Kinase 3 (GRK3) [26]. Prior KORp antibody characterization established that genetic deletion of either GRK3 [27] or prodynorphin [28] blocked stress-induced increases in KORp-immunoreactivity (KORp-IR). Kappa receptor phosphorylation at this site engages arrestin-dependent signaling, which has been associated with aversive and dysphoric qualities of KOR agonists in midbrain circuits [21]. We first confirmed that KOR agonist (U50,488, 10 mg/kg, i.p.) administration significantly increased KORp-IR in PFC (**Figure 1A**) and this increase could be blocked by systemic pre-treatment with the selective KOR antagonist, norBNI (10 mg/kg; **Figure 1B**).

**Figure 1.**
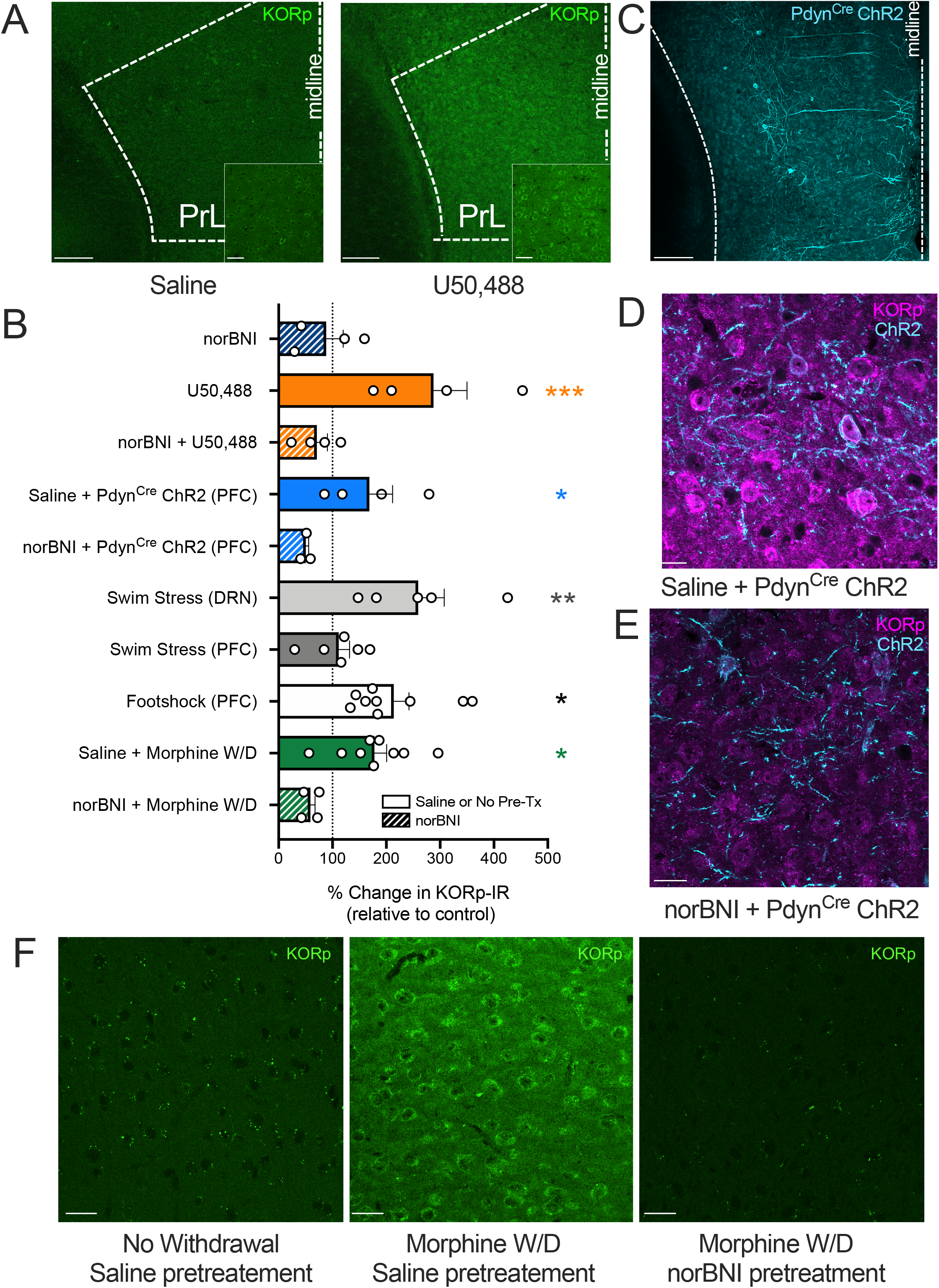
KOR agonists, optogenetic stimulation of PFC dynorphin neurons, or environmental stressors increase KOR phosphorylation in the PFC. **A.** *Pharmacological validation of phospho-KOR antibody*. Representative images (10x, 40x inset) show increased fluorescent signal from phospho-KOR (S369 site) immunoreactivity (KORp-IR) following KOR agonist (U50,488; 10 mg/kg) treatment, compared to saline. Dashed lines show boundaries of the prelimbic prefrontal cortex. Scale bar for 10x image shows 150 *μ*m, scale bar for 40x image (inset) shows 30 *μ*m. **B.** *Conditions for KOR activation in the prefrontal cortex*. All groups were normalized to their corresponding control group. A one-way ANOVA showed an overall significant effect of treatment (F(9,42) = 5.597, p < 0.0001). A Fisher’s LSD posthoc test was used for planned comparisons between groups that were run in the same cohorts and normalized to their own saline or no-stress controls. KOR agonist (10 mg/kg U50,488; n = 4) significantly increased KORp-IR compared to norBNI treatment alone (p = 0.0006). Pre-treatment with norBNI (n = 4) blocked the increase in KORp-IR following U50,488 treatment (p = 0.0002). These groups were normalized to saline-treated controls (n = 4). Optogenetic stimulation (n = 4) of PFC dynorphin neurons (normalized to EYFP control; n = 2) significantly increased KORp IR, which was blocked by norBNI pretreatment prior to optogenetic stimulation (n = 3; p = 0.0468), demonstrating the presence of a local PFC dynorphin/ kappa opioid receptor circuit. Repeated forced swim stress, a stressor known to cause dynorphin release in the brain, significantly increased (p = 0.0024) KORp IR (normalized to no-stress controls n = 5 DRN; n = 6 PFC) in the dorsal raphe nucleus (DRN; n = 6) compared to the prefrontal cortex (PFC; stress n = 6) in the same mice. This showed that swim stress did not cause the release of dynorphin in the prefrontal cortex. In contrast, repeated footshock (0.3 mA shock each min, for 15 min) increased KORp IR in the PFC compared to control mice (one sample t-test; p = 0.0036). Naloxone (1 mg/kg)-precipitated withdrawal (W/D) following chronic experimenter-administered morphine (10 mg/kg, 4-days twice per day, one morphine injection 2 h prior to naloxone on day 5) treatment (normalized to chronic saline treatment n = 8) also increased KORp IR in PFC (n = 9), which was blocked by pretreatment with norBNI (n = 5) when given 24 h prior to 1^st^ morphine injection and 24 h prior to naloxone injection (p = 0.012). Error bars indicate SEM. *p< 0.05; **p < 0.01; ***p<0.0001 **C.** *Representative image for ChR2 expression in PFC dynorphin neurons*. Prodynorphin Cre (pdyn^Cre^) mice were injected in the PFC with Channelrhodopsin-2 (ChR2; AAV1-DIO-ChR2). Image shows ChR2 (cyan) expression in pdyn-Cre^+^ neurons in the PFC. 10X image; Scale bar shows 150 *μ*m. **D.** *Representative image for ChR2 expression and KORp IR in PFC dynorphin neurons with saline pre-treatment*. Image shows 40x image of ChR2 (cyan) and KORp IR (magenta) in mice treated with saline prior to optical stimulation. For optical stimulation, mice were tethered to a fiberoptic patchcord and received 473 nm light (10 mW @ fiber tip) over 30 min (Duty cycle: 5s on @ 20 Hz, 5s off). Within ten min after the end of optical stimulation, brains were perfused for tissue processing. 40X image; scale bar shows 20 *μ*m. **E.** *Representative image for ChR2 and KORp IR in PFC dynorphin neurons with norBNI pretreatment*. Pretreatment with norBNI (10 mg/kg; 24 h prior to stimulation) blocked ChR2-mediated increase in KORp IR. 40X image, scale bar shows 20 *μ*m. **F.** *Representative images for morphine withdrawal (W/D) KORp IR*. Saline treatment after repeated morphine treatment (left) did not produce withdrawal and did not increase KORp IR. Naloxone-precipitated morphine withdrawal increased KORp IR in the PFC during withdrawal, which was blocked by norBNI (10 mg/kg) pre-treatment. 40x image; Scale bars show 20 *μ*m.

We determined that a subset of cortical neurons express dynorphins by injecting *Pdyn*^Cre^ mice [24] with AAV5-DIO-eYFP (**Figure S1A**). We hypothesized that these neurons were likely to release dynorphin locally within the PFC and tested this by optically stimulating the neuronal population and measuring KORp-IR in this region. *Pdyn*^Cre^ mice were injected in the PFC with AAV5-DIO-Channelrhodopsin-2 (ChR2; **Figure 1C**) and >4 weeks later received a 30-min fiberoptic photostimulation period within the PFC (473 nm; 10 mW; 20 Hz; 5s on/5s off), previously shown to evoke dynorphin release in striatum [29]. Local photostimulation significantly increased KORp-IR in the PFC (**Figures 1B-D)**. This increase in KORp-IR was also blocked by pre-treatment with systemic norBNI, indicating that the stimulation protocol evoked dynorphin release in the PFC (**Figures 1B, E)**.

Stress-induced release of the endogenous dynorphin opioid peptides is known to produce both analgesia [30] and aversion [31] in mice, and we next explored behavioral stimuli predicted to physiologically evoke dynorphin release in PFC. Repeated forced swim stress has been shown to stimulate dynorphin release in the dorsal raphe nucleus (DRN), ventral tegmental area, basolateral amygdala, and other brain regions [31; 32]. However, in the present study we found that 2-days of repeated swim stress did not significantly change KORp-IR in the PFC in C57BL/6 male mice, although we did confirm that KORp-IR was significantly increased in DRN in these same mice (**Figures 1B, Figure S1B**). The anatomical selectivity of this stress effect was unexpected. In contrast, repeated footshock, which is known to produce intense cortical activation [33], significantly increased KORp-IR in the PFC (**Figure 1B)**.

Acute abstinence in persons with substance use disorder is profoundly dysphoric and animal models have demonstrated that enhanced dynorphin tone contributes to the aversive state and reinstatement risk during drug withdrawal [8, 34]. To assess whether dynorphin release contributes to the stress responses evident during opioid abstinence, we made mice opioiddependent by daily administration of morphine (10 mg/kg; twice per day for 4 days), then acutely precipitated morphine withdrawal with the opioid receptor antagonist naloxone (1 mg/kg) [35] 2h following a single injection of 10 mg/kg morphine on day 5. Withdrawal behaviors were evoked by naloxone in morphine-dependent mice [36], and some of these behavioral responses (fecal boli) were significantly attenuated by norBNI (**Figure S1C-E**). Naloxone-precipitated morphine withdrawal significantly increased KORp-IR in the PFC (**Figures 1B,F**). The increase in KORp-IR was blocked by norBNI pretreatment prior to morphine administration and withdrawal, demonstrating that dynorphin can be released in the PFC by abstinence-induced stress.

### A genetically encoded fluorescent sensor, kLight1.2a, reveals the dynamics of dynorphin release *in vivo* in the PFC during morphine withdrawal

Immunohistochemistry provides low temporal resolution for measuring the pattern of dynorphin release in the PFC. To achieve higher temporal resolution, we imaged dynorphin release using in vivo fiber photometry and an improved version of the ligand sensor kLight (kLight1.2a; Figure 2A) [37]. We first confirmed that kLight1.2a had the sensitivity and selectivity required. kLight1.2a is based on an inert KOR and a circularly permuted green fluorescent protein (cpGFP) which couples ligand-induced conformational changes of KOR to the fluorescence changes of cpGFP. Neuro2A cells expressing AAV1-hSyn-kLight1.2a responded with a significant increase in fluorescence (ΔF/F) following treatment with the kappa agonists U50,488 (10 μM) or 10 μM dynorphin B (Figure 2B, C). In contrast, treatment with 10 μM morphine did not increase fluorescence. To determine *in vivo* sensitivity, AAV1-hSyn-kLight1.2a was injected into the PFC and after >4 weeks for expression (**Figures 2D, E**), individual mice were injected with saline or U50,488 (1, 5, or 10 mg/kg) on separate days and kLight1.2a fluorescence was monitored by fiber photometry in the PFC as diagrammed (**Figure 2F**). While saline injection did not affect fluorescence (ΔF/F) compared to the pre-injection 5-min baseline period, treatment with U50,488 (1-10 mg/kg) significantly increased fluorescence within 5 min following intraperitoneal administration **(Figure 2G, S1F)**. When mice were pre-treated with a high dose of naloxone (10 mg/kg) sufficient to block KOR [38] 30-min prior to a 10 mg/kg U50,488 injection, there was no significant increase in fluorescence compared to saline. These experiments confirmed that kLight1.2a expressed in the PFC would respond to a systemic KOR agonist.

**Figure 2.**
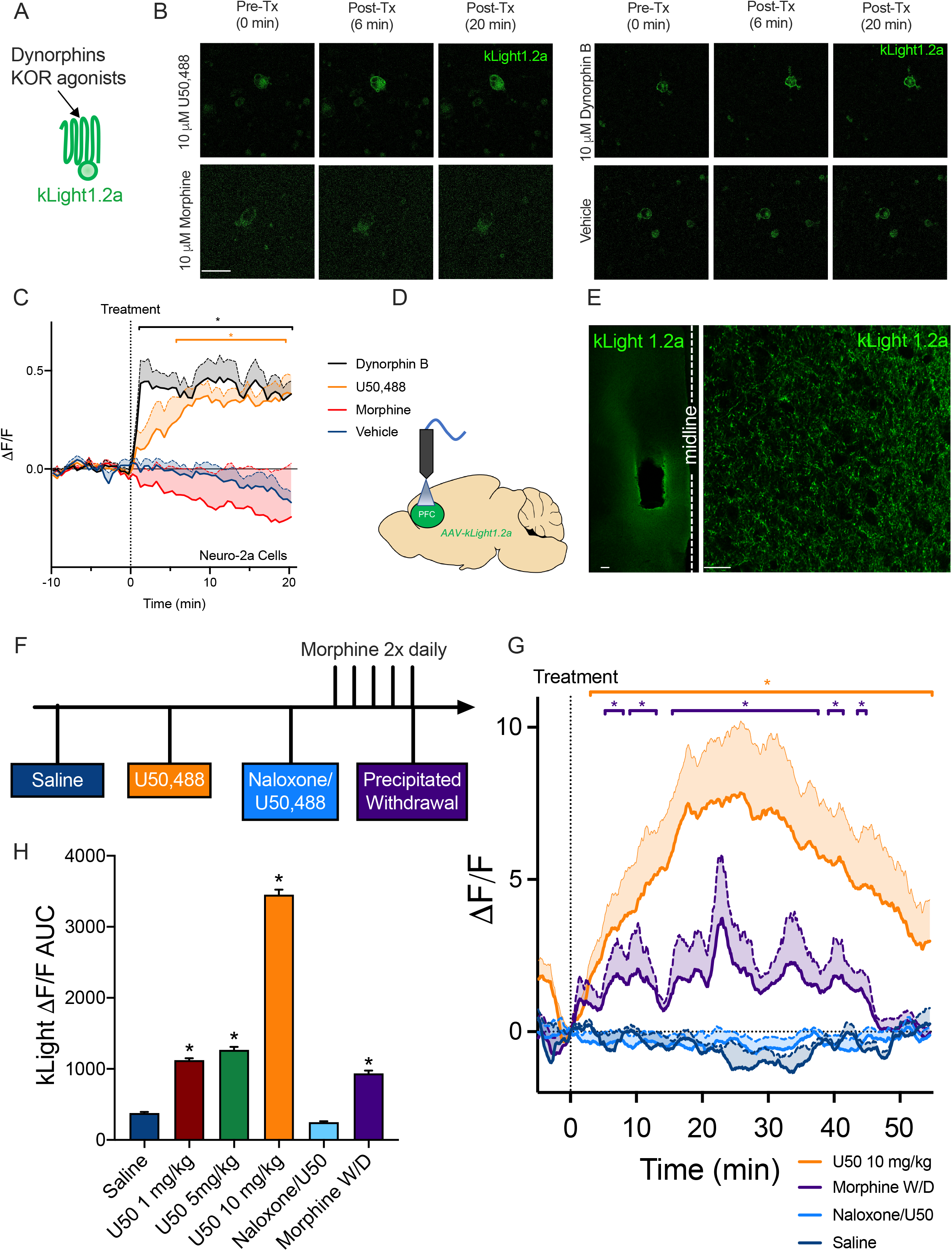
A dynorphin ligand sensor (kLight 1.2a) detects the time course of drug withdrawal-induced dynorphin release in the PFC in vivo. **A.** *kLight 1.2a diagram*. kLight 1.2a is composed of an inert KOR protein fused to a circular green fluorescent protein that responds to conformational changes in KOR when bound to dynorphin or other KOR ligands. **B.** *Representative images of Neuro-2a cells expressing kLight1.2a*. Images show three timepoints, 0.25 min prior to drug treatment (labeled 0), 5.75 (6) and 19.75 (20) min after drug treatment. (Scale bar shows 75 *μ*m) U50,488 (10 μM; upper left) or dynorphin B (10 μM; upper right) treatment increased cell fluorescence. Morphine (10 μM; bottom left) and vehicle treatment did not produce a significant change in fluorescence. **C.** *KOR ligands increase kLight1.2a fluorescence*. U50,488 (n = 6 cells) and dynorphin (n = 4) treatment significantly increased fluorescence (ΔF/F) during the 30-min imaging session starting at 5.75 min (U50,488) and 1.25 min (dynorphin B) following drug treatment. This increase persisted until the end of the imaging session. There was a significant interaction between drug treatment and time (F (180, 1380) = 5.930, p < 0.0001) and Dunnett’s posthoc showed significant (p<0.05) differences between dynorphin and U50,488 compared to vehicle (n = 12). There was no significant effect of morphine treatment (n = 5) on kLight fluorescence. **D.** *kLight1.2a procedure*. KOR^Cre^ mice (n = 5) were unilaterally injected in the PFC with an AAV-kLight1.2a and an optical fiber for photometry was implanted above the injection target region. **E.** *Representative images for kLight1.2a*. kLight1.2a (green) expression was confirmed in mice after recordings. Fiber track for PFC implant visualized in 10x image (left), and 63x (right) image of expression shown on the right. Scale bar for 10x shows 100 *μ*m and scale bar for 63x shows 20 *μ*m. **F.** *Experimental timeline*. kLight1.2a fluorescence was measured using fiber photometry (30 *μ*W at fiber tip; 331 Hz; wavelength 470/405 nm). Following a 5 min baseline recording period, mice were injected with saline or U50,488 (1, 5, and 10 mg/kg) and allowed to freely explore a novel context. Naloxone (10 mg/kg) pretreatment occurred following a baseline period and 30 min later kLight1.2a response to 10 mg/kg U50,488 was recorded. For morphine withdrawal, mice underwent repeated morphine (10 mg/kg) injection procedure, and were tested for baseline fluorescence prior to naloxone (1 mg/kg) precipitated withdrawal. Each mouse received each condition on separate days. **G.** *kLight1.2a detects endogenous dynorphin release in the PFC during morphine withdrawal*. Change in fluorescence (ΔF/F) over a 60-min period is shown (solid line). SEM. is shown above the mean with a filled area. A two-way ANOVA (drug X time) showed a significant interaction between treatment and time (F (3565,14260) = 3.776, p < 0.0001). Dunnett’s posthoc tests showed that U50,488 (10 mg/kg) injection significantly increased kLight fluorescence from 3.3 to 54.5 min compared to saline injection. Pretreatment with the nonselective opioid antagonist naloxone (10 mg/kg) blocked 10 mg/kg U50,488-mediated increases in kLight fluorescence. During the 60-min following 1 mg/kg naloxone administration to precipitate withdrawal in morphine-dependent mice, there was a significant increase in kLight fluorescence from 5.3 to 8.0, 9.0 to 13.0, 15.3 to 37.8, 39.0 to 41.5, and 43.5 to 44.8 min. This showed that kLight detected endogenous dynorphin release within the PFC of mice undergoing morphine withdrawal and that a 10 mg/kg dose of naloxone blocked U50488-mediated increases in kLight fluorescence. **H.** *Dose-dependent effects on kLight fluorescence*. Area under peak values were compared between saline and all other conditions with a one-way ANOVA. There was a significant effect of treatment (F(5,21378) = 561.6, p < 0.0001), and Dunnett’s posthoc showed that saline treatment was significantly different (p < 0.0001) compared to morphine withdrawal or U50,488 (1, 5, and 10 mg/kg). There was no difference between saline treatment and the naloxone/U50,488 group. Error bands above the mean indicate SEM. *p< 0.05

To measure endogenous dynorphin release, we recorded kLight1.2a fluorescence in the PFC during naloxone precipitated withdrawal in morphine dependent mice. At a dose able to block mu but not kappa opioid receptors, naloxone (1 mg/kg) challenge significantly increased fluorescence in the PFC of morphine-treated mice (**Figure 2G)**. There was a significant increase in area under the curve between saline and U50,488 (1, 5, or 10 mg/kg) or during morphine withdrawal (**Figure 2H)**. These results confirmed the KORp imaging data and revealed that dynorphin release could be detected in the prefrontal cortex 5-45 min following the initiation of morphine withdrawal.

### PFC KOR activation disrupts delayed alternation performance via pre- and postsynaptic mechanisms

We hypothesized that activation of the dynorphin/KOR system in the PFC would disrupt cognitive function in a working memory task. In an operant delayed alternation task [17], mice were initially trained to press a retractable lever for a food pellet reward. An alternation contingency was then introduced, during which mice were required to make a response on one retractable lever, wait a specified delay for reintroduction of the levers, and then respond on the alternate lever for reinforcement (**Figure 3A)**. Mice were trained until reaching stable performance with a 10s delay between lever presentations (>20 sessions). Raster plots of performance (**Figure 3B**) show lever presses from a representative mouse early in training (*Line 1*) and following acquisition of the delayed alternation task (*Line 2*). After reaching stable performance in the 10s delay contingency (**Figure S2A**), mice were stereotaxically injected bilaterally in the medial PFC with either artificial cerebrospinal fluid (aCSF) or the long-lasting KOR antagonist norBNI (0.5 μg/0.2 μL aCSF) to locally inactivate KOR in PFC (**Figures S2B,C**). KOR inactivation following norBNI treatment persists for >2 weeks following drug clearance [39] by a molecular mechanism previously described [40]. Following recovery and re-stabilization of performance, mice were treated with either saline (intraperitoneal, IP) or the selective KOR agonist U50,488 (5 mg/kg; IP) immediately before a delayed alternation session. In control mice receiving aCSF in the PFC (PFC aCSF), systemic KOR activation decreased the percentage of correct responses compared to saline treatment (*Line 3*; **Figure 3B**). In mice having norBNI locally administered in the PFC (PFC norBNI), systemic KOR agonist administration produced no significant degradation in performance during early (0-15 min) or late phases (45-60 min) of the delayed alternation session (*Line 4*; **Figure 3B)**. In both the early phase and the late phase of the session, KOR activation disrupted performance in PFC of aCSF injected mice by decreasing the percent of correct responses. There was also a KOR-mediated decrease in total responses during the early phase, but no significant difference in total response number during the late phase (**Figures 3C-F).** PFC norBNI administration blocked the response suppressive effect of PFC KOR activation on early phase responding.

**Figure 3.**
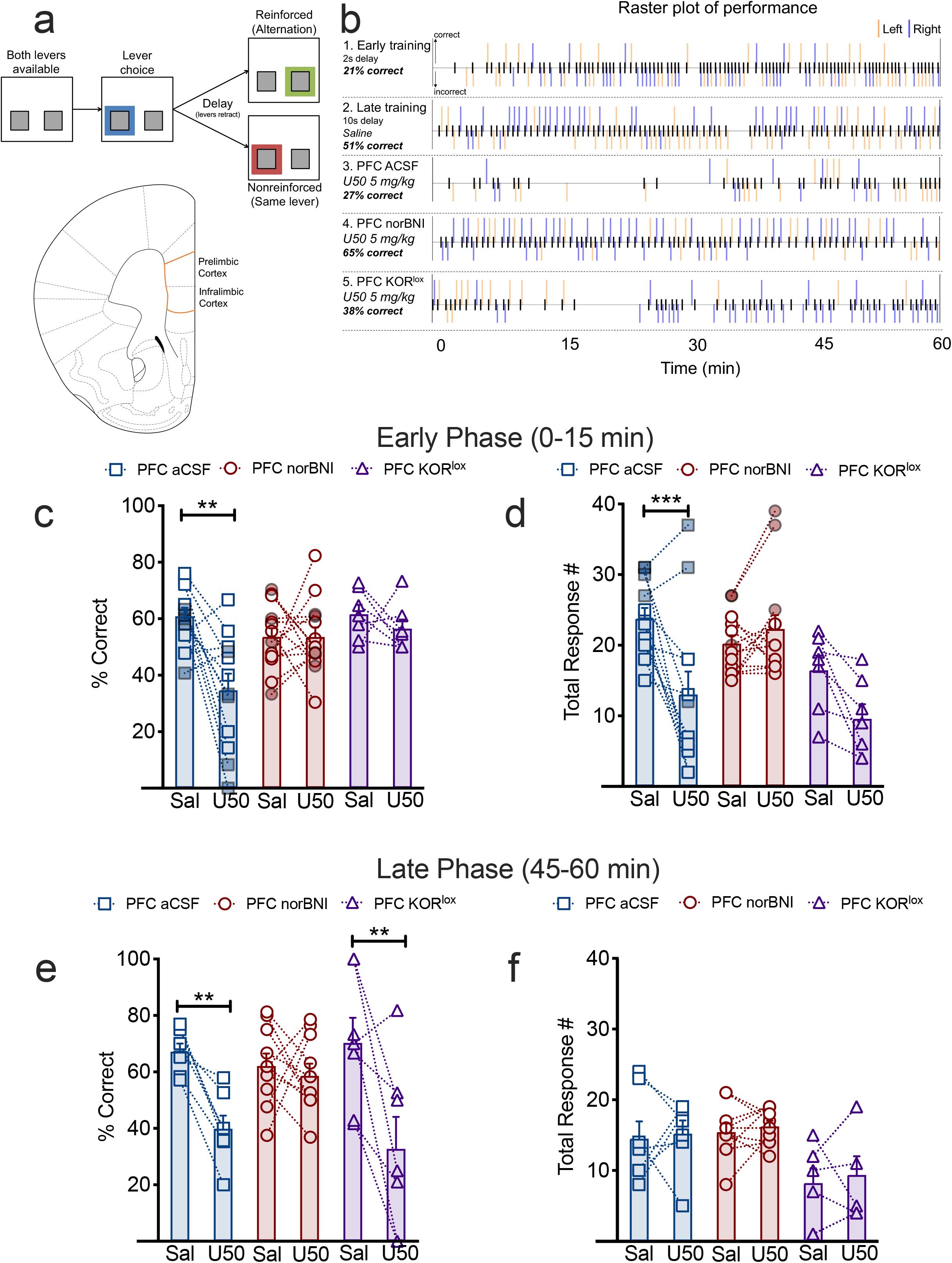
KOR activation disrupts delayed alternation performance in early and late phases of the test session. **A.** *Delayed alternation procedure*. C57BL/6 male mice were food restricted and trained to complete lever alternations for reinforcement (detailed description in Methods). At the start of a trial, both levers were extended. Lever choice initiated a 10s delay period during which both levers remained retracted and unavailable. When levers were re-inserted, mice chose the alternate lever for food pellet reinforcement. Choosing the same lever as the previous response was recorded as an incorrect response and was not reinforced. After a 20s intertrial interval, levers were reinserted to allow mice to start the next trial. The diagram of a hemi-coronal section outlines the PFC region (infralimbic and prelimbic cortex) targeted for injection; all placements were confirmed postmortem. **B.** *Raster plots of performance of a representative mouse during a 60-min trial session*. Ticks (black) on the center line show trial initiating responses. Ticks above the center line show correct (reinforced) responses and ticks below show incorrect (nonreinforced) responses. Lever choice is shown by orange (left lever) or blue (right lever) ticks. Line 1 shows performance of a PFC aCSF mouse early in delayed alternation training (with a 2s delay). Mice rarely alternated levers early in training, leading to a significant number of response errors. Raster line 2 shows improved performance in the same mouse after several training sessions, and saline injection (IP) prior to the delayed alternation session did not significantly alter performance in the task. Raster line 3 shows that treatment with the KOR agonist U50,488 (5 mg/kg, IP) in the same mouse decreased the percent of correct responses and decreased total response number during the delayed alternation session. Line 4 shows that bilateral local injection of the KOR antagonist (norBNI; 0.5 *μ*g/ 0.2 *μ*L) in the PFC blocked the disruptive effects of systemically administered U50,488 on both total response number and percent correct responses. Line 5 shows that U50,488 treatment decreased the correct responses in mice with postsynaptic KOR deletion (KOR^lox/lox^ mice injected with 0.2 *μ*L AAV5-Cre-eYFP bilaterally in the PFC; PFC KOR^lox^) during the late phase of the session, but not during the early phase of the session. **C.** *Postsynaptic KORs are required for early phase agonist-mediated decreases in correct response % in delayed alternation*. After training for 3 wk, aCSF (n = 12) or norBNI (n = 14) was bilaterally injected into PFC (3-5 d prior to testing), and the first fifteen minutes of performance were compared between saline and KOR agonist (U50,488) treatment days. There was a significant interaction between group and agonist treatment F(2,30) = 5.600, p = 0.0086. A Sidak’s post-hoc test showed a significant difference between U50 and Saline treatment in the PFC aCSF mice (p = 0.0003), but not in PFC norBNI or PFC KOR^lox^ mice (n = 7). Gray borders and filled symbols indicate mice that received 30 min, rather than 60 min, training and testing sessions. **D.** *Systemic KOR agonist significantly decreases total responses in PFC aCSF mice*. Total number of alternation responses was significantly decreased in PFC aCSF mice during the first 15 min of the session (Drug X Group Interaction: F(2,30) = 8.582, p = 0.0011; Sidak’s post-hoc: p = 0.002). There was a nonsignificant trend towards a decrease in response number in the PFC KOR^lox^ group (p = 0.0921). Gray borders and filled symbols indicate mice that received 30 min, rather than 60 min, training and testing sessions. **E.** *Postsynaptic KOR deletion does not block late phase KOR-mediated decreases in delayed alternation performance*. KOR activation (PFC aCSF n = 7; norBNI n = 10; KOR^lox^ n = 7) significantly decreased percent correct in the delayed alternation task (F(2,21) = 4.761, p = 0.0197) during the last 15 min of the session (late phase; 45-60 min) in PFC aCSF (Sidak’s; p = 0.0171) and PFC KOR^lox^ (p = 0.0011) mice. **F.** *KOR activation did not affect total response number during the late phase of a delayed alternation session*. There was no significant effect of drug on total response number following KOR activation. Dashed lines connect responses of individual mice receiving saline 1 d prior to the U50,488 trial session. Error bars indicate SEM.*p<0.05, **p < 0.01; ***p<0.0001

To assess the role of postsynaptic KOR in PFC, we injected AAV5-DIO-Cre in floxed KOR (KOR^lox/lox^) mice [21] to selectively excise KOR in PFC neurons (PFC KOR^lox^; **Figure S2D)**. KOR-mediated decreases in percent correct in the early phase were not evident in PFC KOR^lox^ mice, whereas disruptions in the late phase were unaffected (**Figures 3C-F; S2E)**. In contrast, local norBNI blocked disruptions at both phases. This pattern of behavioral disruptions suggests that KORs in postsynaptic PFC neurons are required for the early phase of cognitive disruptions and presynaptic PFC KORs mediate the later phase of cognitive disruption.

### Dynorphin release in the PFC disrupts delayed alternation performance

We tested whether opioid withdrawal-induced dynorphin release would alter delayed alternation performance. Mice were initially trained to stable performance in the task with a 10s delay as described above. Following a baseline testing session, mice received four days of 10 mg/kg morphine twice daily. On the fifth day, mice were treated with morphine 2 h before the test session, then received naloxone immediately prior to entry into the operant chamber (**Figure 4A**). There was not a significant effect of precipitated withdrawal on early phase task performance (**Figure 4B**). However, during the later phase of the delayed alternation session (30-45 min) mice showed a significant impairment in performance. This impairment was not blocked by postsynaptic deletion of KOR in PFC (**Figure 4C; Figure S3A-F**), but was blocked by local norBNI injection in the PFC.

**Figure 4.**
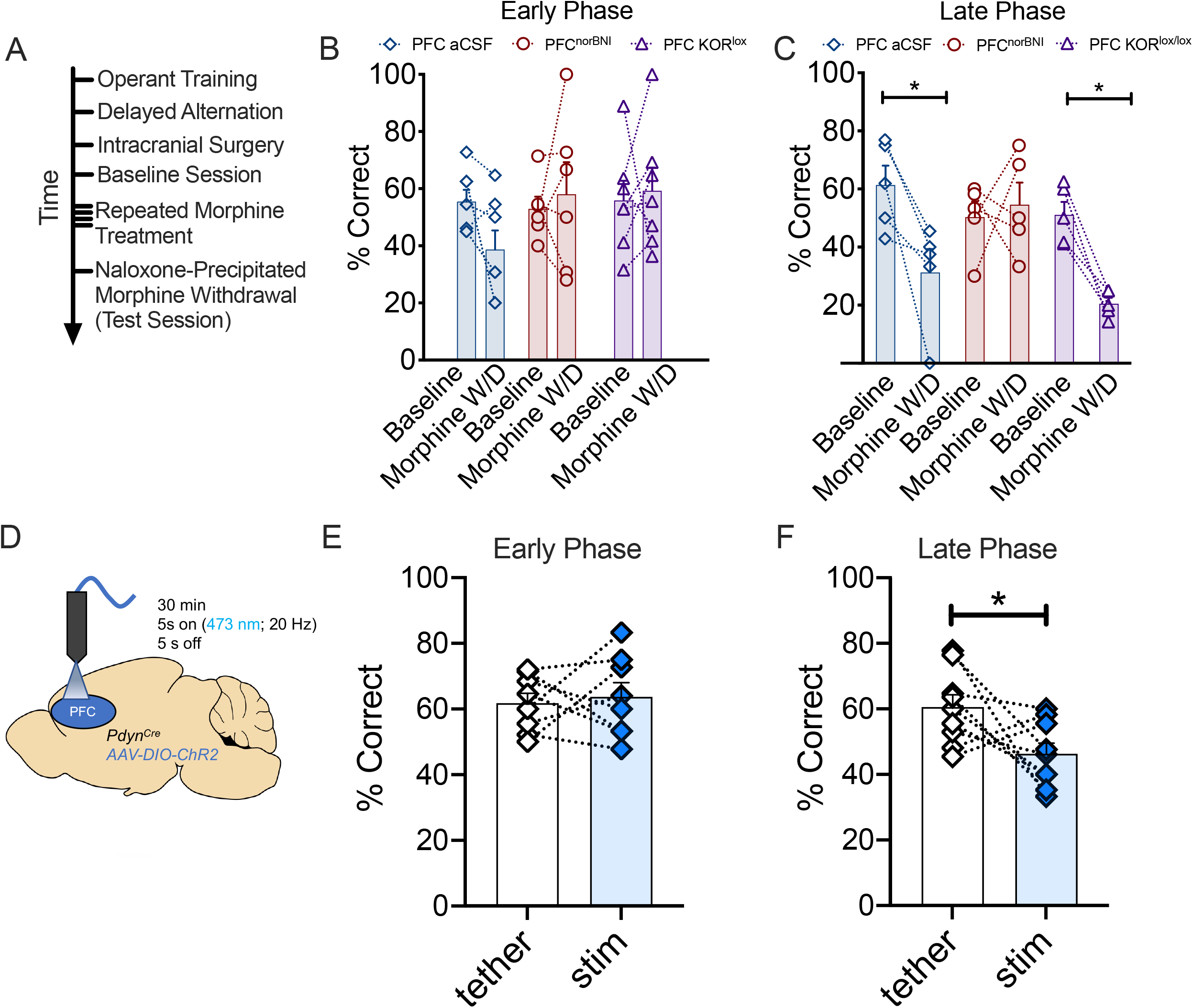
Dynorphin release in the PFC disrupts delayed alternation performance. **A.** *Experimental timeline*. Mice received training in delayed alternation until reaching stable performance, then were intracranially injected with aCSF or norBNI. Floxed KOR mice were injected prior to training. Following recovery, mouse performance in a baseline delayed alternation session was assesed. Mice then received morphine (10 mg/kg) for four days, and on the fifth day received naloxone (1 mg/kg) prior to entry into the operant chamber. **B.** There was no significant effect of morphine withdrawal on performance during the early phase of the task in the aCSF (n = 7), norBNI (n = 6), or floxed KOR (n = 7) groups. **C.** *Morphine withdrawal disrupts delayed alternation performance after naloxone treatment*. In the 30-45 min window following naloxone treatment, there was a significant decrease in percent correct responding during a delayed alternation session in the aCSF group (n = 5), and the disruptive effect was not blocked by postsynaptic KOR deletion (n = 5). However, local norBNI pretreatment (n = 5) in the PFC blocked the disruptive effect of morphine withdrawal on delayed alternation performance. (Main effect of drug; F(2,12 = 4.628; p = 0.0324); Sidak’s posthoc compared to baseline: PFC aCSF(p = 0.0211), PFC norBNI (p = 0.959), KOR^lox^ (p = 0.0191). **D.** *PFC pdyn^Cre^ optical stimulation procedure*. Pdyn^Cre^ mice were injected with ChR2 in the PFC and were trained in delayed alternation. On baseline and test days, mice were tethered to an optical patchcord for 30-min prior to a delayed alternation session and received either no stimulation (tether) or optical stimulation (stim) on separate days. **E.** *Optical stimulation of PFC pdyn^Cre^ neurons does not disrupt early phase delayed alternation*. Optical stimulation had no significant effect on percent correct in delayed alternation in pdyn^Cre^ mice (n = 5) during the early phase of the test session. **F.** *Optical stimulation of PFC pdyn^Cre^ neurons disrupts late phase delayed alternation performance*. Compared to a tether session, photostimulation significantly decreased the percent of correct responses 45-60 min after stimulation (t8 = 2.33, p = 0.048). Error bars indicate SEM. *p< 0.05.

We then tested whether optical stimulation of PFC dynorphin neurons that increased KORp-IR also significantly disrupted delayed alternation performance **(Figures 4D-F)**. For a baseline day, mice were tethered to the optical patchcord for 30 min then performance in the task was determined. On the test day, mice received optical stimulation (30-min session as described for **Figure 1B-D**) then were placed in the operant chamber. There was no difference between optically stimulated or tethered mice during the early phase of the session (**Figure 4E; Figure S3G,H**). However, optical stimulation significantly decreased percent correct responding during the late phase of the task **(Figure 4F)**.

## Discussion

The principal conclusion from this study is that the dynorphin / kappa opioid receptor system functions in the mouse medial prefrontal cortex to control cognitive performance in a working memory task. Both systemic KOR agonist treatment and evoked release of dynorphins activated KORs to reduce response accuracy. The specific contribution of post-synaptic KORs was tested by AAV-Cre induced excision in floxed KOR mice, which showed that KOR activation in cortical neurons reduced response number and accuracy in the early phase of the operant task. Dynorphin release in the PFC elicited by morphine withdrawal or optogenetic stimulation of PFC pdyn^Cre^ neurons also disrupted delayed alternation performance during the session. Our studies strengthen the preclinical basis for investigating KOR antagonists to decrease stress-induced cognitive dysfunction.

We show that aversive stimuli (e.g. footshock stress, drug withdrawal) will cause dynorphin release in the prefrontal cortex. In humans, during the first two weeks of abstinence from opioids, there are significant deficits in multiple working memory domains, in addition to deficits in executive function that can persist for months in opioid dependent individuals [41]. Disrupted cognitive function may contribute to alterations in mood, stress reactivity, and reward processing observed in early opioid abstinence. KOR antagonists may be useful for decreasing some of these symptoms and could promote long-term abstinence by decreasing the disruptive effects of stress on behavior and cortical substrates. Cognitive dysfunction and altered cortical function are also common symptoms of schizophrenia, and some evidence suggests a modulatory role of the dynorphin/KOR system in this disorder. Elevated dynorphin concentrations in cerebrospinal fluid have been reported in individuals with schizophrenia and are associated with poorer treatment outcomes [42, 43]. In double-blind clinical trials, nonselective opioid antagonists reduced auditory hallucinations [44], and KOR antagonism was sufficient to decrease some psychotic symptoms [45]. While still preliminary, these provocative findings suggest that selective KOR antagonists might have also therapeutic utility in the adjunctive treatment of psychosis [46].

Pharmacological KOR activation in the PFC is known to produce anxiety [47] and aversion [48] and KOR/dynorphin activation in other brain areas can also alter cognition by disrupting attention [49-51] or behavioral inhibition (23, 52). KOR activation has immediate and remote effects on behavior [53, 54], and we observed multiple phases of response disruptions during delayed alternation test sessions. We have previously shown that KOR activation produces perseverative responses during the later phase of an operant test session in a behavioral inhibition task [23]. Although KOR activation can significantly suppress responding during operant sessions [49-51], these effects were not evident when KOR activation was blocked in the PFC. Our data indicate that there is a component of the behavioral response being mediated by the prefrontal cortex that contributes to the prolonged suppression of responding caused by systemic KOR activation. We speculate that the initial period of behavioral suppression may be related to the dissociative or hallucinogenic effects of KOR activation, and the late phase error responses may reflect a perseverative mode of responding, leading to decreased lever switching or suppressed ability to inhibit responding. KOR activation likely contributes to disruptions in behavior over relatively long periods of time by producing complex patterns of circuit inhibition and excitation through recruitment of particular intracellular signaling elements and engagement of associated homeostatic responses.

Based on the wide pre- and postsynaptic distribution of KORs and dynorphins in the PFC [15, 24, 47, 48], it is likely that KOR in the PFC has actions in many different circuits and this coordinated disruption of multiple nodes of activity could lead to the loss of cognitive control following stress. Single cell gene expression studies in cortical regions in mice have suggested diverse clusters of cells that are enriched in KOR and prodynorphin expression [55]. KOR activation in presynaptic terminals in the PFC decreases the local release of dopamine from ventral tegmental area neurons [47] and glutamate from basolateral amygdala neurons [48]. We aimed to dissociate pre- and postsynaptic contributions of KOR activation in the prefrontal cortex and found that virally mediated excision of KORs from the PFC blocked disruptions during the early phase of the task. Although adenoviral strategies can lead to some retrograde transport that may delete KOR from select inputs into the prefrontal cortex, we expect that the bulk of the KOR deletion in our experiments is likely to occur in prefrontocortical cells. We observed that KOR agonist-mediated disruptions were blocked by postsynaptic KOR deletion, however withdrawal-mediated disruptions in behavior were not. Together, these data suggest that presynaptic KOR effects contribute to dynorphin-mediated changes in behavior during opioid withdrawal.

Clinical trials with KOR antagonists have shown promising results for further development, but short-acting KOR antagonists will only be effective if used during periods of time in which dynorphin is released. The ligand sensor kLight1.2a enables the measurement of dynorphin release dynamics in the brain and can reveal the types of behavioral or environmental events that are associated with increased dynorphin activity. Opioid withdrawal and abstinence have previously been shown to drive changes in the dynorphin/KOR system [56], but direct evidence for this relationship relied on techniques with low temporal precision, such as immunohistochemistry. In humans, binge psychostimulant use has been associated with increased dynorphin release in the prefrontal cortex [57], and chronic opioid use is likely to produce similar increases in PFC dynorphin release in humans. Using kLight1.2a, we demonstrated that naloxone-precipitated morphine withdrawal increased dynorphin release in the prefrontal cortex of mice, however the cell populations involved in this effect are unknown. Our immunohistochemical studies show that dynorphin is released from *pdyn*^Cre^ PFC neurons following optogenetic stimulation and the same stimulation paradigm produced disruptions in delayed alternation behavior, suggesting that PFC dynorphin neuron activation could contribute to dysfunction in working memory. However, *pdyn*^Cre^ neurons are likely to release multiple signaling molecules following optical stimulation, and further characterization of the subpopulations of dynorphin neurons in the PFC is required to determine the signaling pathways through which withdrawal-mediated effects might occur.

Overall, these studies determined spatial and temporal aspects of stress-induced cognitive dysfunction. Increased dynorphin release in the prefrontal cortex following stress disrupted cognition, suggesting that KOR antagonists could be useful for treating cognitive symptoms in stress-vulnerable illnesses like substance use disorders. Our studies indicate that effective clinical translation of kappa therapeutics will require further analysis of the contributions of the KOR/dynorphin system to cognition in animal models and humans.

## Funding and Disclosure

Funding sources: P50-MH106428 (CC), P30-DA048736 (CC), R01-DA030074 (CC), R21-MH108839 (BL), T32-DA07278 (CC and AA). The authors have nothing to disclose.

## Acknowledgements

We thank Mackenzie M. Andrews, Zeena M. G. Rivera, and Juliana Chase for assistance with data collection and data analysis for this manuscript.

## Author contributions

ADA, SMC, and SSS conducted the experiments, ADA and CC wrote the manuscript, BAW generated software for data extraction and analysis, GOM, KM, and SSS generated reagents for experiments, ADA, BBL, and CC designed the experiments, CC and LT secured funding.

## Supplemental Information

### Detailed Methods

#### Cell Culture

Neuro2a cells were plated in 12 well dishes and transduced with AAV1-Syn-kLight1.2a. Cells were plated on chambered coverslips (ibidi GmbH, Gräfelfing, DE) 24-48 hours after transduction. Cell media was replaced 6 h prior to the experiment, which took place 3-4 days after transduction. Cells were imaged on a Leica SP8X confocal microscope using LASX software at 20× with a 2× digital zoom, maintaining temperature in the imaged chambers at 34.9 °C (S.D., ±0.2 °C). Two treatment conditions were imaged in parallel for each experiment, tracking two fields per condition using the “hold focus” and “mark and track” functions, with scanning intervals of 30 s. kLight1.2a (excitation with WLL 488 laser at 10%, emission with HyD sensor at 498-–619 with 0.5% gating and smart gain of 100%) was imaged with 512 × 512 resolution, 600 speed with bidirectional scanning, and live averaging of 2. Twenty frames (10 min) were measured before the first drug treatment; cells were then treated as described using 10x drug stocks in media. Data were analyzed in ImageJ 1.52e. Fluorescence was quantified by drawing a box around each cell which encompassed the cell in all frames, and then quantifying the average pixel intensity of that ROI in each frame. Fluorescence at each time point was subtracted from the average fluorescence during the baseline period, and divided by average baseline fluorescence and to produce ΔF/F values. To remove linear drift, cells were detrended using linear regression of ΔF/F during the baseline period. Cells which detached, went out of focus, or moved out of the field of view were excluded. Four cells (2 vehicle, 1 morphine, 1 dynorphin) were excluded based on significant linear fluorescent drift over the imaging session.

#### Genotyping

All transgenic mice were genotyped using Transnetyx (Cordova, TN, USA) genotyping services. Prodynorphin-IRES-Cre mice were genotyped by DNA isolated from tail tissue obtained from weanling mice (21-28 days of age), and PCR screening was performed for the presence of Cre recombinase. For KOR^lox^ mice, the following primers were used for PCR screening: Forward Primer: CACTTTTAAACATGGAGTAGGGTGATG; Reverse Primer: GGCCGCATAACTTCGTATAGCATA; Reporter: CCGGTGCTTCTGTGTATC.

#### Stereotaxic injection

Mice were anesthetized with isoflurane and mounted on a stereotaxic alignment system (Model 1900 David Kopf Instruments, CA, USA). A sterile ophthalmic ointment (Puralube; Dechra, KS, USA) was applied to the eyes to prevent drying. A 30 gauge needle Neuros Syringe (Hamilton, NV, USA) was lowered bilaterally for PFC (A/P = +1.75 mm; M/L = ±0.3 mm; D/V -2.8 mm) injections and artificial cerebrospinal fluid (aCSF) or 0.5 μg norBNI in 0.2 μL aCSF was injected bilaterally. Stereotaxic injections of norBNI occurred in mice that had been trained for stable performance in the delayed alternation task. After injection, the needle was kept at the injection site for 5 additional minutes before removal. To confirm targeting, a subset of mice was injected with carboxylate-modified orange (540/560 nm) fluorescent 0.04 *μ*m microspheres (Invitrogen; Carlsbad, CA) lightly coating the tip of the Hamilton syringe. Animals were sutured with 5–0 polypropylene sutures (Sharpoint, PA, USA) and allowed to recover for 3-5 days before behavioral testing began. For KOR^lox^ mice, 0.2 μL of AAV5-Cre (Addgene, MA, USA) was injected bilaterally. For pdyn^Cre^ mice, 0.2 μL of AAV1-DIO-hChR2(H134R)-EYFP-WPRE (Addgene) was injected bilaterally for behavioral experiments or unilaterally for a subset of immunohistochemistry experiments. After viral infusion, the needle was kept at the injection site for at least 5 min before removal. A chronically implantable fiberoptic cannula (photometry: 400/430 core, 0.57 NA (kLight1.2a); (optogenetic stimulation: 200/240 core, 0.22 NA; Doric Lenses, Quebec, CA) was placed 0.3 mm dorsal to the viral injection site and dental cement (Stoelting, Wood Dale, IL, USA) was used to secure the cannula to the skull. Viral infusions occurred at least 4-6 weeks before pharmacological studies, optical stimulation, or fiber photometry.

#### Operant Conditioning Apparatus

Standard mouse operant chambers (model ENV-300, Med Associates, VT, USA) equipped with fans and housed in sound-attenuating chambers were used to measure delayed alternation in mice. Each chamber was equipped with a house light, a sound generator, two ultra-sensitive retractable levers and a food receptacle (magazine). A house light illuminated the box during lever access periods. A computer equipped with the MED-PC IV program (Med Associates) controlled the apparatus and recorded lever presses and head entries into the magazine.

#### Fixed Ratio

Training sessions occurred over 60 min. At the start of each training session, the ventilation fan was turned on and one retractable lever (left or right) was designated as the active lever. Prior to the first training session, a 20-mg food pellet was placed in the food magazine and the active lever to encourage nose poke and magazine entry behaviors. Lever press lead to automated delivery of food into the food receptacle. Mice were trained on a fixed ratio 1 (FR1) schedule, with each single lever press leading to the delivery of a single food pellet. The active lever was alternated during training to discourage lever bias. Mice were trained on an FR1 schedule for 3-8 d, until reaching stable high responding for food pellets (>30 responses per 60-min session). Approximately 10% of mice did not acquire fixed ratio responding over 10 sessions and were removed from the experiment.

#### Delayed Alternation

Following acquisition of FR1, mice were introduced to an alternation contingency, where a mouse is required to make a response on one retractable lever, wait a specified delay for re-insertion of the levers, and then respond on the alternate lever. Mice typically acquire this task within ~3 weeks. The wait period between lever retraction and insertion was varied to control the difficulty level of the task. Mice were trained until reaching stable performance (>3 days of over 40% correct in a 2s delay then increased to 5s delay, then 10s delay). Approximately 5% of mice were unable to achieve stability in 2s training and were excluded from further training. Levers remained inserted into the chamber until receiving a response. After stable performance was achieved in the 10s delay, mice were injected in the PFC with either artificial cerebrospinal fluid (aCSF) or the long-lasting (~3 weeks) KOR antagonist, norBNI. A subset of mice in the PFC aCSF and PFC norBNI groups received 30 min training and testing sessions, but there were no significant differences in performance between 30- and 60-min groups, so data from these groups were pooled for analysis of early session disruptions. In KOR^lox^ and pdyn^Cre^ mice, viral injection occurred prior to the start of the delayed alternation training. Mice were treated with vehicle (saline) or drug (U50,488) immediately before a delayed alternation session on consecutive days. Mice also received naloxone (1 mg/kg) to precipitate morphine withdrawal immediately before a delayed alternation test session. Between treatments, mice were given multiple (>3) delayed alternation sessions to ensure stable baseline performance (>40%) in the task.

#### Intracardiac perfusions and antigen retrieval

Mice were anesthetized with pentobarbital (Beuthanasia-D) and intracardially perfused with room temperature phosphate-buffered saline (PBS) and chilled 10% formalin. Thereafter, brains were stored overnight in 10% formalin. Brains were sectioned into 5 mm width sections and placed in a small basket in PBS (85° C - 90° C for three min). The brains were agitated every thirty seconds. Immediately after three minutes, the brains were removed from the warmed PBS and immersed in room temperature PBS. Brains were then put in 20% sucrose for long term storage.

#### Immunohistochemistry

Prefrontal cortex slices were sectioned at 40 *μ*m, then washed in PBS before being placed in blocking solution (PBS containing 5% normal goat serum and 0.3% Triton X-100). PFC slices were incubated in a rabbit anti-KORp antibody (1:25 dilution), mouse anti-Cre (1:500; MAB 1320; MilliporeSigma, Burlington, MA, USA), or chicken anti-GFP (1:3000; ab13970; Abcam, Cambridge, UK) solution diluted in blocking buffer (Detailed protocol in Lemos et al. [1] for 72 h on a shaker in a cold room (4°C). KORp peptide was generated by Biomatik (Wilmington, DE, USA), and conjugated with mKLH to increase immunogenicity [2]. Rabbits were immunized with the mKLH-conjugated KORP peptide; following an initial immunization and at least 5 boosters, antisera were harvested and affinity purified (R & R Research; Stanwood, WA, USA). Slices were then washed in PBS and incubated with an AlexaFluor 488 or 555 goat anti-rabbit (A11008; A32732), 488 goat anti-chicken (A11039), or 488 goat anti-mouse (A28175) secondary antibody (1:500 dilution; ThermoFisher Scienitific, Waltham, MA, USA) for two hours covered on the shaker. After 2 h, the slices were washed in PBS and mounted on Superfrost Plus slides with Vectashield hardset mounting media and imaged at a Leica SP8X Confocal Microscope.

#### Morphine Withdrawal

Mice were injected twice daily (9:00AM and 5:00PM) with morphine (10 mg/kg) for four days. On the fifth day, two hours following the AM morphine injection, mice were intraperitoneally injected with 1mg/kg naloxone and were perfused 30 minutes after injection. This dose of naloxone antagonizes mu opioid receptors, while allowing kappa opioid receptor activation [3]. Mice were video-recorded in this 30 min period and behavioral withdrawal data were quantified by an experimenter blind to treatment assignments. For delayed alternation, KOR^lox^ or control mice that previously received U50,488 or nalfurafine were used after restabilization in delayed alternation to test the effect of morphine withdrawal on behavior. Delayed alternation testing occurred prior to chronic morphine injections as a baseline and naloxone treatment immediately preceded placement into the operant chamber to precipitate morphine withdrawal during the test session.

#### Footshock

Mice were placed inside a MED PC operant chamber equipped with scrambled shock floors (ENV-414SA; Med Associates), mice received one 0.3 mA footshock per minute for 15 min.

#### Repeated Forced Swim Stress

To induce stress, mice were exposed to a modified forced-swim test as previously described [4]. Briefly, the modified-Porsolt forced-swim paradigm used a 2-day procedure in which mice swam in 30 °C water in a 5L opaque beaker for 15 min the first day, and then 6 min during each of four trials on the second day (6 min interval between each swim) without the opportunity to escape. Mice receiving repeated swim stress were tested in a delayed alternation session prior to swim stress on Day 1 and immediately following the last swim stress on Day 2.

#### Optogenetic stimulation

A small nestlet was placed under the head of the mouse prior to connecting the incoming fiberoptic patchcord to the indwelling fiberoptic cannula. Mice were placed into a novel cage with fresh bedding and allowed to freely explore with patchcord attached. On tether days, mice received no light stimulation and were placed into the delayed alternation chamber after 30 min of tether habituation. On optical stimulation days, mice received 473 nm light source (10 mW incoming laser power; OEM Laser, Midvale, UT) controlled through a waveform generator (Grass Instruments). Stimulation parameters (5s of 20 Hz (10 ms pulse) laser on, 5s laser off; 30 min session) were selected based on Al-Hasani et al. [5] showing optically elicited dynorphin with 20 Hz stimulation. Following stimulation, mice were perfused or placed into a delayed alternation test session. Mice (n = 5) were tested twice in delayed alternation with each condition (tether or stim) and data were analyzed within subjects. One mouse had to be removed due to headcap loss.

#### Fiber Photometry

We used a real-time signal processor (RZ5P; Tucker-Davis Technologies) connected to Synapse Software (Fiber Photometry) to set frequency of light stimulation and record input from photodetectors. The RZ5P was connected to a light emitting diode (LED) driver (Doric Lenses) that controlled the power of a 470/405 nm (kLight1.2a) Doric LED. The LED was attached with a low autofluorescence patchcord (400/430) to a Fluorescent MiniCube (Doric Lenses) with dichroic mirrors. Optical patchcords connected the integrated Mini Cube/photodetector (ilFMC4; Doric Lenses) with a pigtailed rotary joint (FRJ; Doric Lenses) that allowed free animal movement during data collection. Prior to photometry sessions, patchcords were bleached with light for at least 4 h to minimize autofluorescence. Power of the LED at the fiber tip was set to ~30 μW and was tested prior to the start of each session. Signals were collected at a sampling frequency of 1017 Hz. Each of the sessions were downsampled by a factor of 100 and normalized to a five-minute baseline period in the beginning of the recording. The sessions were then smoothed using a moving average filter (100s window) to remove high frequency noise and detrended to remove linear drift. The isosbestic channel (405 nm) was fitted to the 470 nm channel using a least-squares method and subtracted to remove motion artifacts. Each session started with a 5 min baseline recording period prior to pharmacological experiments to calculate fluorescent change from baseline (ΔF/F; Change in fluorescence from baseline fluorescence/baseline fluorescence). For in vivo kLight1.2a experiments, n = 5 mice were run with all conditions on separate days (Saline; U50,488 10 mg/kg; 10 mg/kg naloxone/10 mg/kg U50,488; 1 mg/kg naloxone precipitated morphine withdrawal, 1 mg/kg U50,488, 5 mg/kg U50,488) with each treatment separated by > 1 week.

---

**S1.**
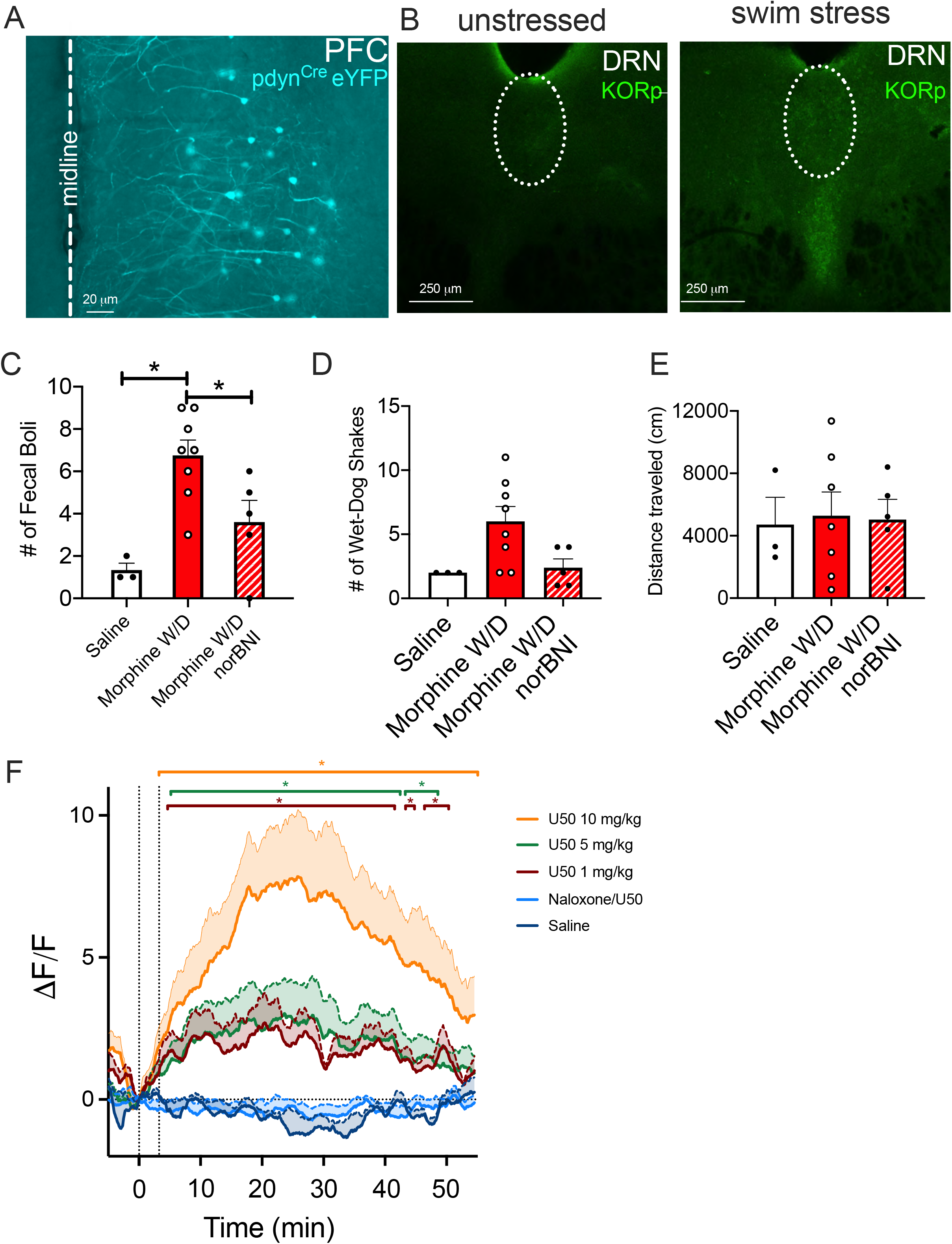
Supplement 1 Legend. **A.** Pdyn^Cre^ injected with eYFP (cyan) in the PFC. **B.** Dorsal raphe nucleus (DRN) KORp representative images. KORp IR was analyzed within the DRN and compared between stressed and unstressed groups. **C.** Naloxone precipitated morphine with low doses of morphine produced a significant elevation in the number of fecal boli observed in mice and no significant difference in **(D.)** wet dog shakes or **(E.)** distance travelled. **F.** kLight1.2a dose in vivo response data is shown. Overall statistical analysis for this experiment is shown in Figure 2 Legend, with all groups included. Compared to vehicle, 10 mg/kg U50,488 significantly increased kLight fluorescence from 3.25 to 54.5 min, 5 mg/kg U50,488 significantly increased kLight fluorescence from 5.2 to 42.4 min and 43.3 to 48.5 min, and 1 mg/kg U50,488 significantly increased kLight fluorescence from 4.4 to 41.5 min, 43.5 to 44.7 min, and 46.5 to 50.0 min.

**S2.**
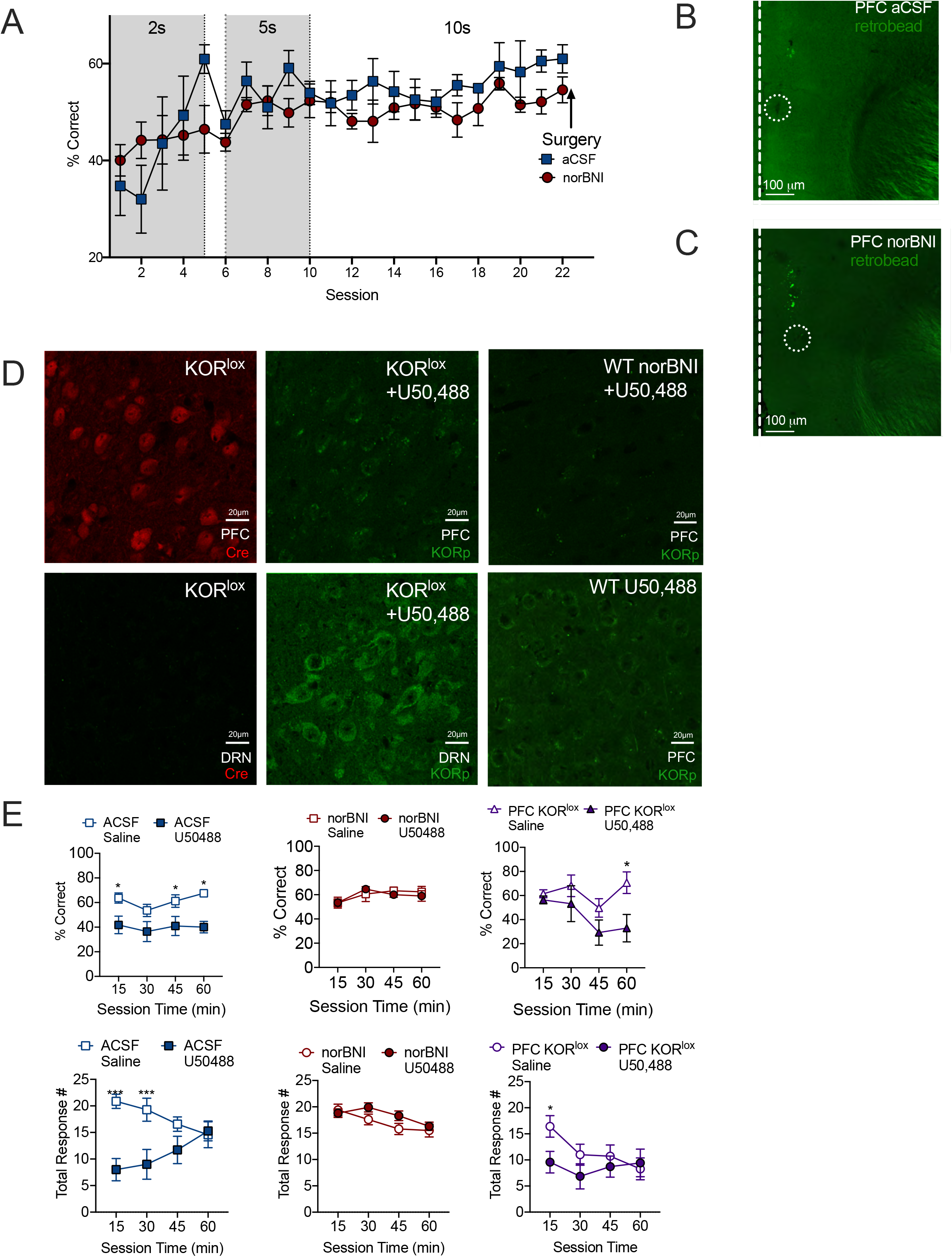
Supplement 2 Legend. **A.** *Delayed alternation acquisition*. Mice were trained for approximately five sessions in 2s and 5s delays until performance reached over 50% on average. Then, all mice were trained in 10s delay until performance was consistently over 50% prior to microinjections into the PFC with aCSF or norBNI. Mice were randomly assigned to a treatment condition. **B.** *Microinjection targeting images for aCSF **(B)** or norBNI* **(C).** Mice were injected several weeks before perfusion with a single microinjection rather than implanted cannula. Due to this procedure, damage to the PFC was minimal. To aid in visualizing the injections, needle tips were dipped in fluorescent retrobeads prior to injection, and traces of fluorescence can be observed in the PFC in both groups. **D.** KOR^lox^ mice injected with AAV-Cre in the PFC were treated with U50,488 prior to tissue collection to visualize KORp immunoreactivity. Immunohistochemistry with anti-Cre antibody (red) shows Cre expression in KOR^lox^ mice (upper left), and KORp IR measurement (green) confirms the lack of KORp IR fluorescence in the PFC (top center). In the same mouse, KORp IR was present in the dorsal raphe nucleus (DRN; lower center), and there was no Cre expression (lower left), as expected. In a wildtype mouse injected with U50,488 (10 mg/kg; lower right), KORp immunofluorescence appeared increased and localized within cells compared to KOR^lox^ mice or WT mice pretreated with norBNI (10 mg/kg) prior to U50,488 treatment (upper right). These images demonstrate that norBNI pre-treatment completely blocks KORp fluorescence in the PFC, whereas postsynaptic deletion of KOR in the PFC of KOR^lox^ mice leaves some remaining KORp IR fluorescence which appears to localize differently than the cellular staining observed in intact WT U50,488 mice. **E.** Time course for each group (PFC aCSF, PFC norBNI, PFC KOR^lox^) across the 60-min session with percent correct in top row, and total response number in lower row. *p < 0.05. Error bars indicate SEM.

**S3.**
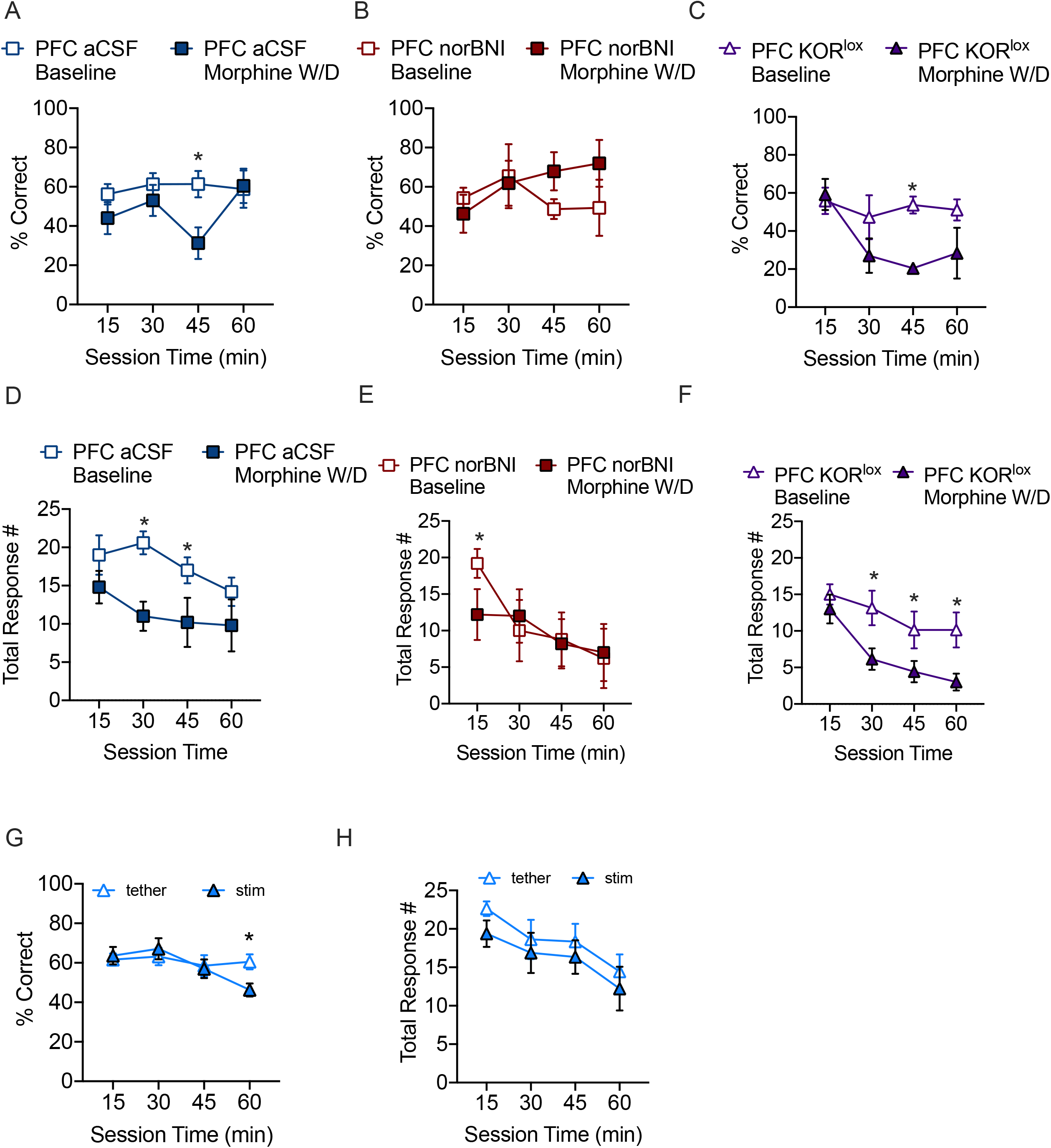
Supplement 3 Legend. **A-C**. Time course of percent correct from delayed alternation shown for morphine withdrawal in each group (**A**) PFC aCSF, **(B)** norBNI, or **(C)** mice with KOR deletion in the prefrontal cortex (KOR^lox/lox^) is shown in this figure. Sidak’s post-hoc test results shown. **D-F.** Total alternation responses from **(D)** PFC aCSF, **(E)** norBNI, or **(F)** KOR^lox/lox^ groups. **E-F.** Time course of **(E)** percent correct and **(F)** total alternations in prodynorphin^Cre^ mice following stimulation of Channelrhodopsin in PFC dynorphin neurons. *p<0.05. Error bars indicate SEM.

## Notes

### Competing Interest Statement

The authors have declared no competing interest.

